# Bacterial diversity of chronic middle ear lesions revealed by high-throughput sequencing

**DOI:** 10.1101/2021.02.10.430714

**Authors:** Xingzhi Gu, Xiuqing Cheng, Tuoheti Abulajiang, Xiaoban Huang

## Abstract

Chronic otitis media is a common middle ear disease in otolaryngology and head and neck surgery. Bacterial infection is considered the main cause of disease, but relying on conventional bacterial cultures can be problematic for identification of specific pathogens. Current research suggests that bacteria in microbial communities can only be identified by rDNA sequencing of bacteria.This prospective study utilized broad-range PCR amplification of 16S rRNA genes with clone analysis to compare bacterial diversity in lesions from 6 patients with chronic suppurative otitis media (CSOM) and 10 patients with cholesteatoma of middle ear lesions. Bacteria were analyzed at the phylum, order, family, genus, and species levels. Bacterial species abundance and species diversity were greater in cholesteatoma of the middle ear lesions than in CSOM lesions. At all bacterial taxonomic levels, the epithelial tissue of middle ear cholesteatoma was complex in terms of bacterial diversity, covering a large number of Gram-positive and Gram-negative bacteria, likely related to bacterial microbiome formation. In contrast, bacteriology of the CSOM lesions was relatively simple at all taxonomic levels, with all sequences characterized as belonging to Gram-negative bacteria. These results suggest that persistent infection of middle ear cholesteatoma may be a microbial flora disorder, which is related to conditional pathogenic bacteria, rather than a single bacterial infectious disease. Findings from the study may have implications in the selection of antimicrobial agents for the treatment of chronic otitis media.

## INTRODUCTION

Chronic otitis media, a common disease in otorhinolaryngology, is usually caused by inflammatory lesions in the middle ear and/or mastoid cavity, and is characterized by ear discharge (otorrhea) and hearing loss. Some patients experience serious complications that can be life-threatening (1, 2). Clinically, chronic otitis media usually refers to chronic suppurative otitis media (CSOM) and middle ear cholesteatoma. Bacterial infection is considered to be the main pathogenic factor of chronic otitis media (3).

Bacterial culture of secretions from the middle or outer ear canal is often used to confirm the presence of pathogenic bacteria, and various pathogenic bacteria, even methicillin-resistant *Staphylococcus aureus*, have been obtained through conventional bacterial culture (4).

Sensitive antimicrobial agents based on the culture results have been used to treat patients with success, but there remains some cases where the treatment is ineffective (5, 6). Subsequent studies confirmed that bacterial biofilm formation leading to resistance to antimicrobial agents may be one reason for the persistence of symptoms in some patients (7–10). Bacterial culture results are affected by many factors including the inability of some bacteria to grow on standard medium and restrictions on specimen collection. Thus, in some cases it is difficult to successfully identify and treat the causative pathogen (11).

In recent years, there have been numerous reports on molecular biology techniques for the study of bacteria of otitis media. No bacteria were isolated from middle ear secretions of adults and children by conventional bacterial culture methods, but gene sequencing resulted in the isolation of various bacterial sequences, including the phylum Thunb and the genus Grapevine (12). Neeff et al. demonstrated the use of molecular biological techniques to analyze CSOM and healthy human middle ear mucosa, which can more accurately evaluate the information of microbial communities in the middle ear or mastoid mucosa (13). However, there are limited reports on the distribution of microbes in different types of chronic otitis media. In the current study, the molecular technique of 16S rRNA gene amplicon sequencing was employed to study microbial communities in CSOM and cholesteatoma of middle ear (CME) diseased tissues. Differences in microbial species abundance, diversity, and at the levels of phylum, class, order, family, genus, and species were compared. These analyses provide a basis for future studies on the characterization of bacteria at the molecular level in chronic otitis media.

## MATERIALS AND METHODS

### Patient information

All patients in this study had chronic otitis media and received surgical treatment in the otolaryngology department of Xinjiang Uygur Autonomous Region People’s Hospital from January 2017 to June 2019. Among the 16 eligible patients, six cases were chronic suppurative otitis media and 10 were middle ear cholesteatoma. None of the patients used antibiotic ear drops or systemic antibiotics prior to surgery, but according to the requirement of preventive use of antibiotics, all patients received intravenous cephalosporins 30 min ~ 2 h before surgery. Patients were not associated with basic metabolic diseases such as diabetes or rheumatism, and did not have fungal infections of the external auditory canal.

The study was approved by the Ethics Committee of the People’s Hospital of Xinjiang Uygur Autonomous Region (Xinjiang District Hospital Ethics Committee 2015054).

All patients were admitted to Xinjiang Uygur Autonomous Region People’s Hospital for surgery. The operation mode of canal wall up [CWU] tympanic forming or canal wall down [CWD] tympanic forming was selected according to the disease severity of the patient. After opening the tympanic sinus and mastoid cavity during the operation, the middle ear lesion tissue was retained (after cleaning the matrix of cholesteatoma, the cholesteatom aepithelium and granulation tissue of cholesteatoma were retained). Following collection of pathological tissues, the accompanying blood and floats were washed with normal saline, immediately placed into a specimen tube, frozen in liquid nitrogen, and then stored at −80°.

#### Absolute quantification of 16S rRNA amplicon sequencing

Absolute quantification of 16S rRNA amplicon sequencing was performed by Genesky Biotechnologies Inc. (Shanghal, 201315, China). Briefly, total genomic DNA was extracted using a Fast DNA SPIN Kit for soil (MP Biommedicals, Santa Ana, CA, USA) according to the manufacturer’s instructions. The integrity of the genomic DNA was assessed through agarose gel electrophoresis, and the concentration and purity of genomic DNA were measured using a NanoDrop 2000 spectrophotometer and Qubit 3. 0 fluorometer.

GC content were artificially synthesized. Then, an appropriate proportion of spike-ins mixture with known gradient copy numbers were added to the sample DNA. The V3-V4 hypervariable regions of the 16s rRNA gene and spike-ins were amplified with the primers xxxF (5’-CCTACGGGNGGCWGCAG-3’) and xxxR (5’-GACTAACHVGGGTATCTAATCC-3’) and then sequenced using an Illumina NovaSeq 6000 sequencer.

#### Sequence data processing and analysis

Raw read sequences were processed in QIME2 [14]. The adaptor and primer sequences were trimmed using the cutadapt plugin, and the DADA2 plugin was used for quality control and to identify amplicon sequence variants (ASVs) [15]. Taxonomic assignments of representative ASVs were performed with a confidence threshold of 0.8 by a pre-trained Naive Bayes classifier that was trained on the Greengenes database (version 13.8). The spike-in sequences were then identified, and reads were counted. A standard curve for each sample was generated based on the read counts versus spike-in copy number, and the absolute copy number of ASVs in each sample was calculated by using the read counts of the corresponding ASV. Since the spike-in sequence is not a component of the sample flora, the spike-in sequence was removed in the subsequent analysis [16].

## RESULTS

### Analysis of species abundance and diversity

High-throughput 16SrRNA gene sequencing technology was used to sequence samples from 16 patients with CSOM or cholesteatoma. A total of 4,672,171 raw sequence reads and 3,415,184 optimized sequence reads were obtained from the 16 patients with middle ear lesions (6 CSOM cases, 10 cholesteatoma cases). The average sequence length from the total of 1,243,460,816 bases was 364.10 bp.

The number of sequences obtained from each patient group were 582,102 and 1,485,436 for CSOM and cholesteatoma, respectively. Chao 1, ACE, Shannon, and Simpson indices indicated there were no differences in α-diversity between the two disease groups. However, coverage was significantly different between the two groups (P=0.022478), illustrating that the coverage of microbial communities in cholesteatoma samples was higher than that in the CSOM group.

### Taxonomic analysis of pathological tissue flora from patients with CSOM or cholesteatoma

A total of 833 operational taxonomic units (OTU) were subjected to taxonomic classification, and 28 phyla, 50 classes, 63 orders, 128 families, 264 genera, and 232 species were identified and further analyzed.

Samples from different disease groups have certain characteristics and commonalities. Based on the OTU abundance table, the unique OTU, and common OTU in each group were screened and visualized. The total number of detected OTU was 838, comprising 788 OTU detected in cholesteatoma pathological tissues, 230 in CSOM pathological tissues, and 180 OTU common to both groups (FIG 2).

**FIG 1.**
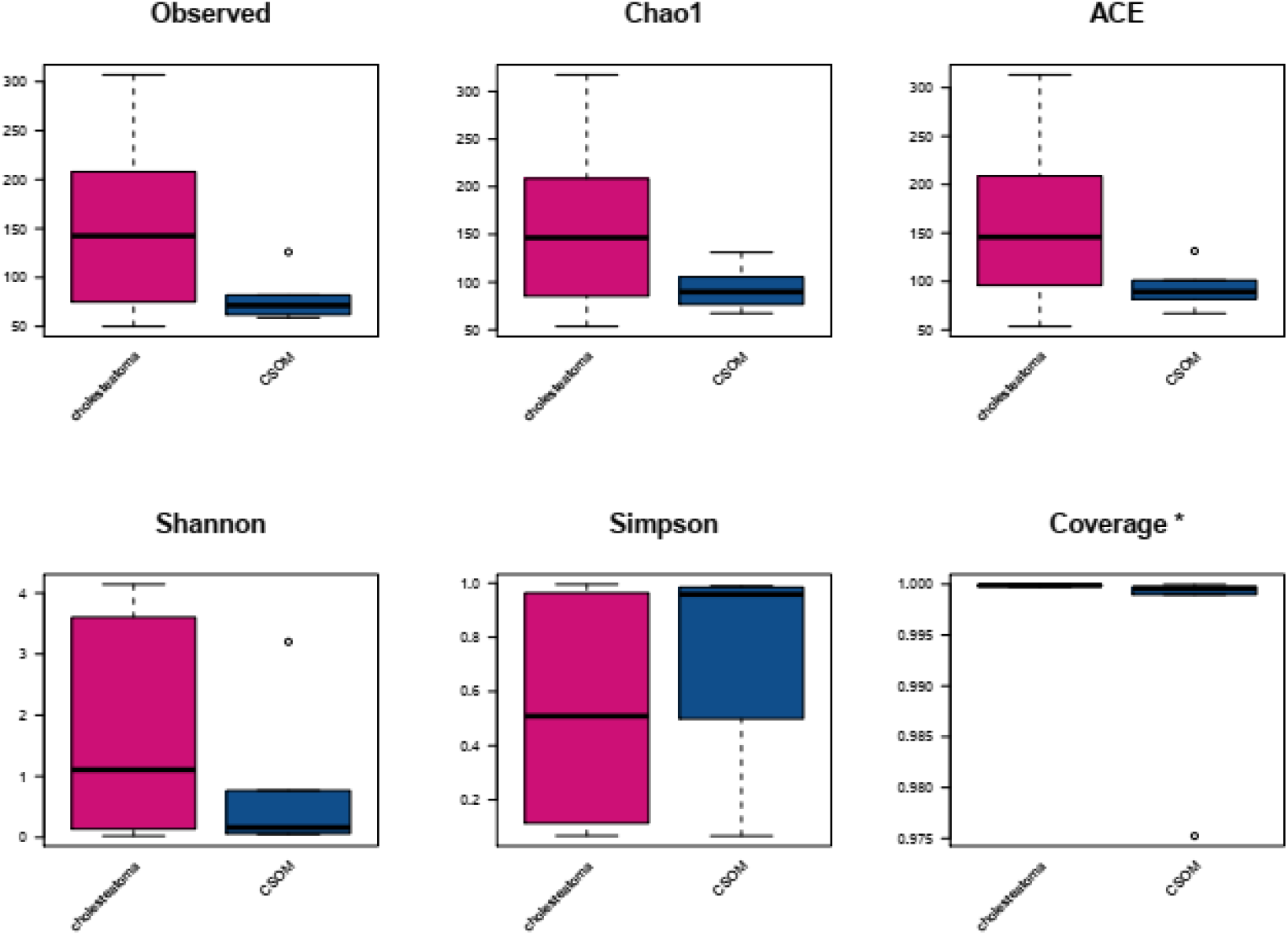
Box diagrams of α-diversity. A: Observed; B: Chao 1 index, P=0.093407; C:ACE, P=0.093407; D: Shannon index, P=0.3131868; E: Simpson index, P=0.562188; F: Coverage, P=0.022478. Asterisk indicates statistical significance at P≤0.05.

**FIG 2.**
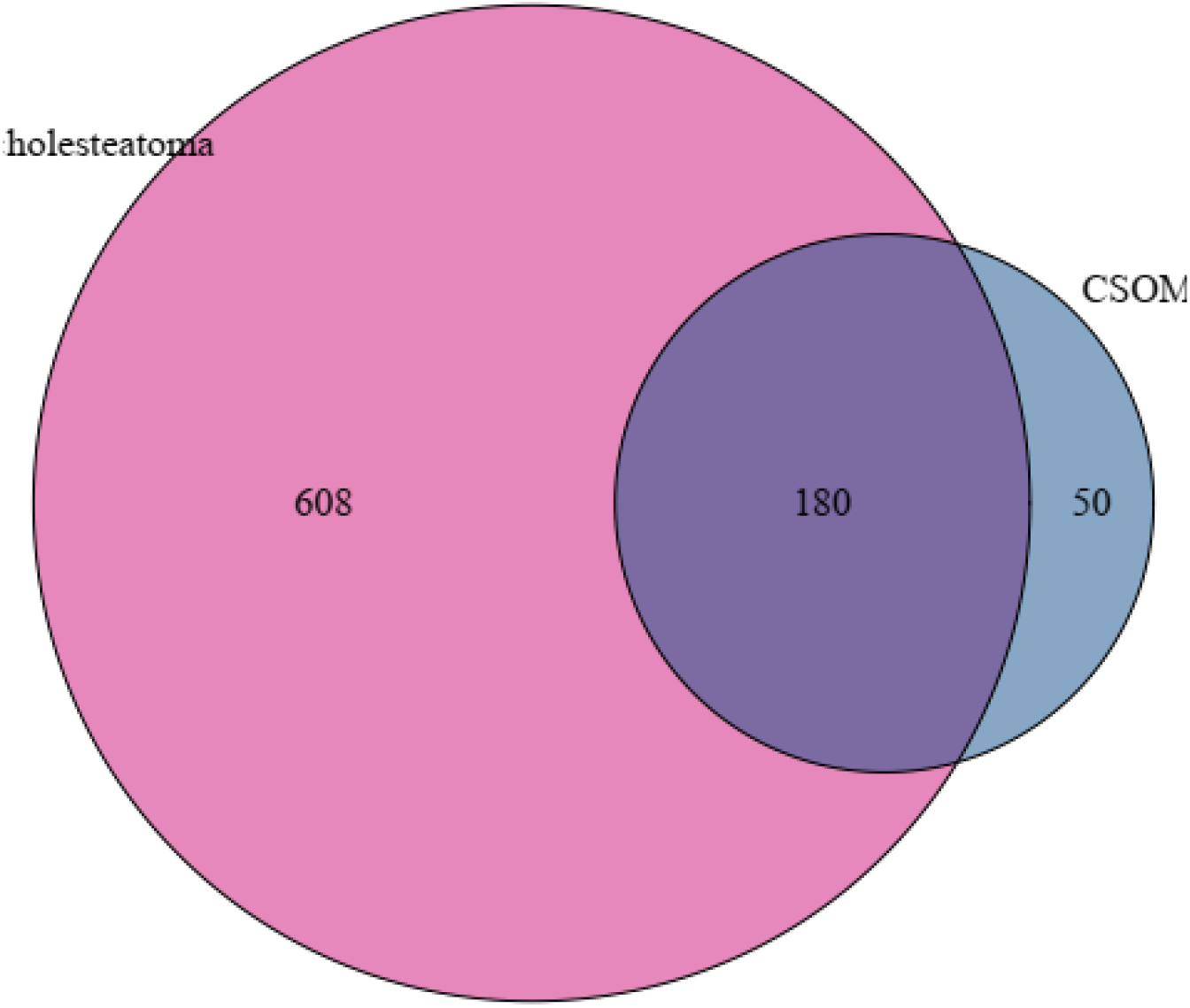
Venn diagram of OTUs of CSOM and cholesteatoma lesions

### Phylum

Bacterial community analysis at the phylum level revealed that the community flora of CSOM tissues was simpler than that of Cholesteatoma samples. The phylum *Proteus (Proteobacteria)* was detected in both groups. Statistically, detection of Proteus indicates this phylum is a major part of CSOM rt (99.46%, P=0.000321). And *Pseudomonas aeruginosa*, *Escherichia coli* and so on in the door of Proteus are also our common Gram-staining-negative pathogens. The microbial community of Cholesteatoma samples was more complex. Aside from the detection of Proteus (*Proteobacteria*) (35.77%), statistically significant phyla of the Cholesteatoma samples were thikum (*Firmicutes*) (44.21%, P=0.001071) and *Actinomycetes* (*Actinobacteria*) (16.66%, P=0.032464). These two phyla encompass the vast majority of Gram-positive bacteria (FIG 3).

**FIG 3.**
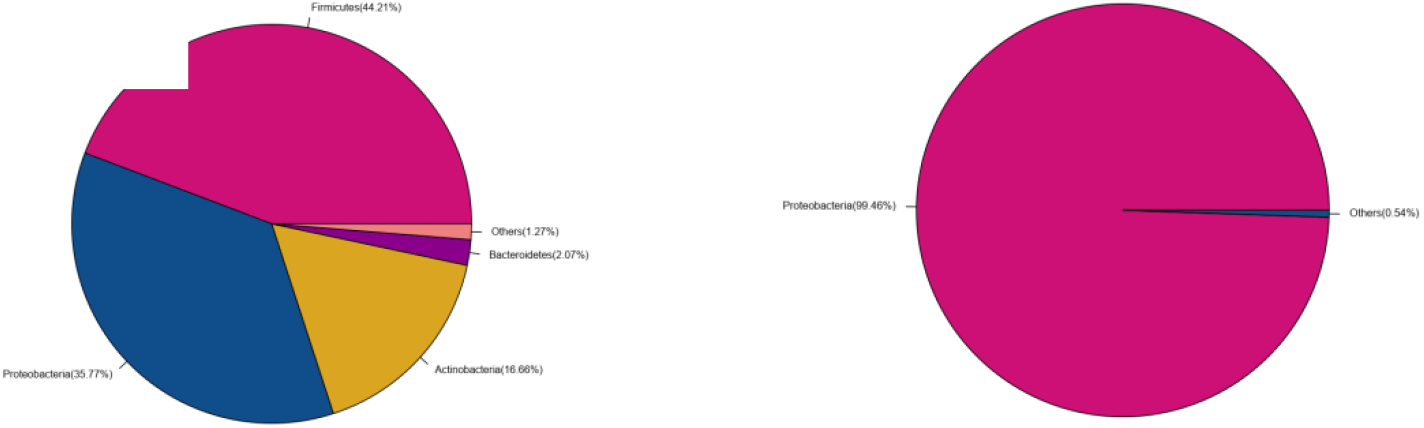
Pie charts showing phylum breakdown of bacterial communities obtained from CSOM and cholesteatoma pathological tissues.

### Class

Analysis of bacterial sequences at the class level showed statistical differences between the two groups (CSOM and cholesteatoma) in the classes *Gammaproteobacteria* γ-Proteus (P=0.03), *Bacilli* (P=0.01034), *Clostridia* (P=01034), and *Bacteroidia* (P=0.041) (FIG 4). In general, the measured species of the cholesteatoma lesions were higher than those of the CSOM group, indicating that the abundance of microbes was greater. In the CSOM group, *Gammaproteobacteria* γ- and β- Proteus (*Betaproteobacteria*) were predominantly detected, indicating that the number of microbial species present at the class level is relatively small in this group (FIG 5).

**FIG 4.**
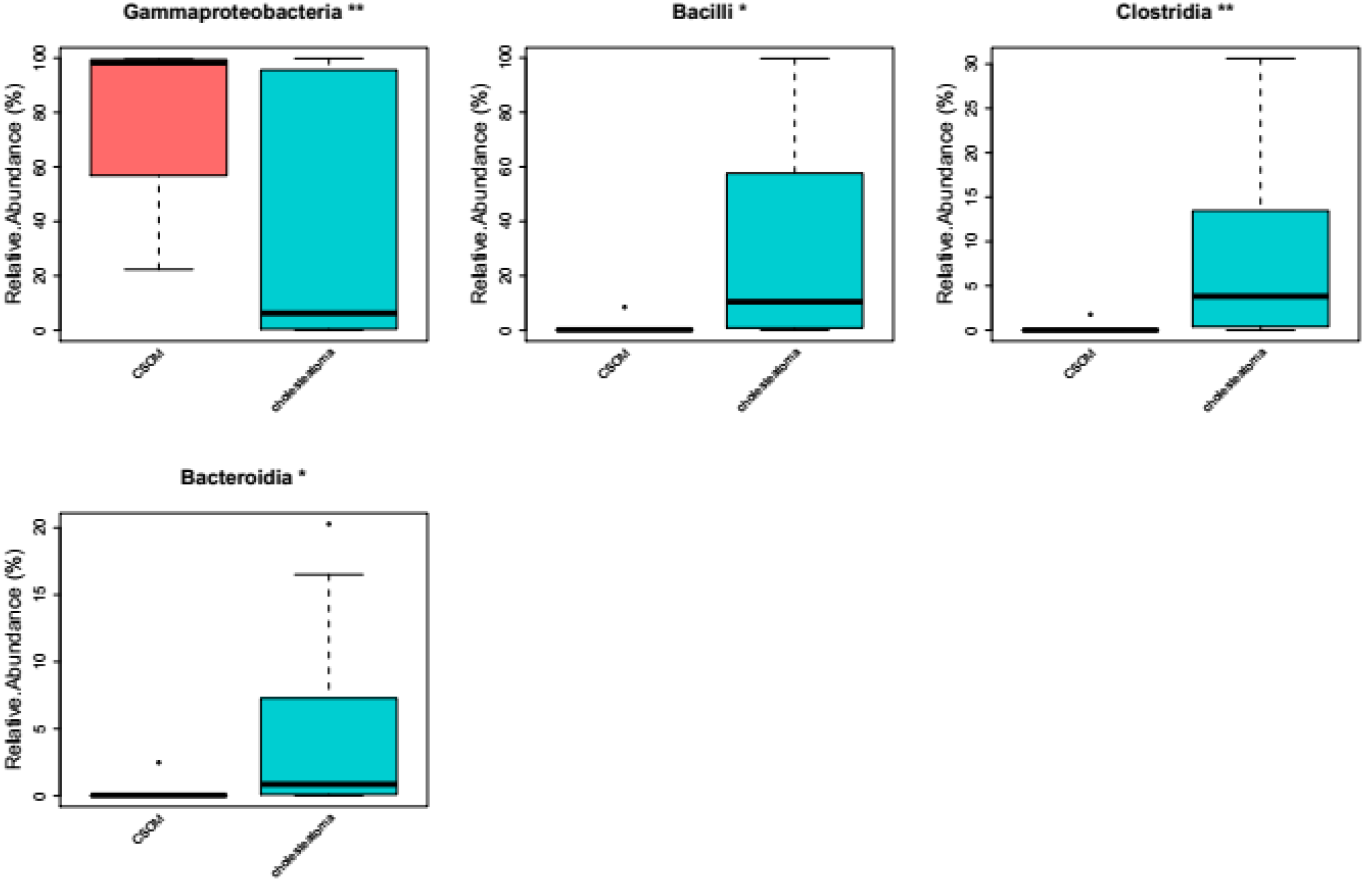
Class level differences in bacterial community of CSOM and cholesteatoma pathological tissues

**FIG 5.**
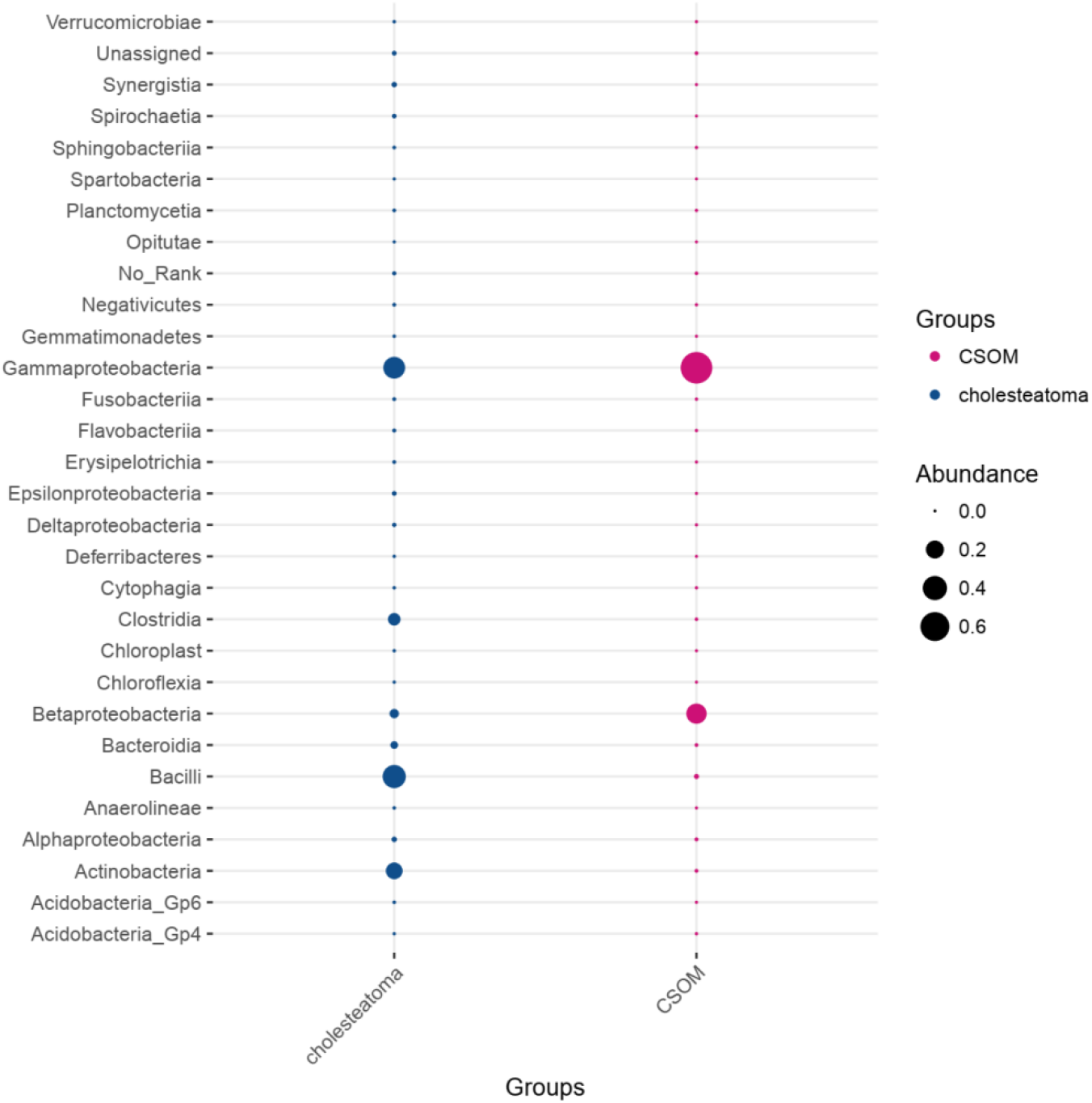
Bubble diagram of class level differences in bacterial communities of CSOM and cholesteatoma pathological tissues

### ORDER

At the order level, *Pseudomonadales* and *Burkholderiales* have an absolute advantage in CSOM, with *Pseudomonadales* being statistically significant (P=0.00129). *Pseudomonas aeruginosa* strain PAO was identified in the bacterial species map at the sample sequencing level. In the cholesteatoma group, the statistically significant orders were *Bacillales* (P=0.011887), *Clostridiales* (P=0.011887), and *Bacteroidales* (P=0.002532) (FIG6). Description of the bacterial communities at the order level also suggests that cholesteatoma is more complex in bacterial diversity (FIG 7).

**FIG 6.**
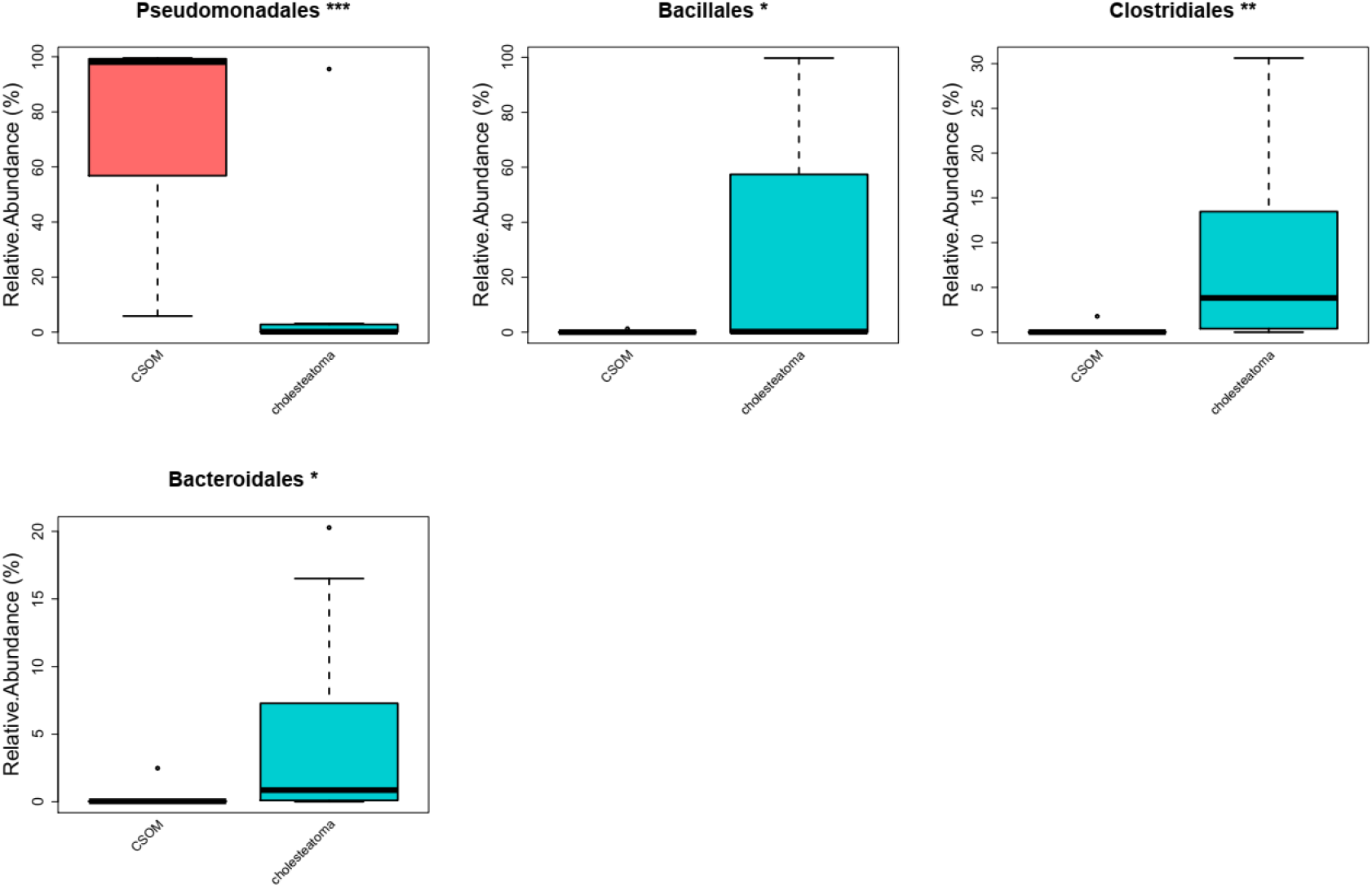
Order level differences in bacterial communities of CSOM and cholesteatoma pathological tissues

**FIG 7.**
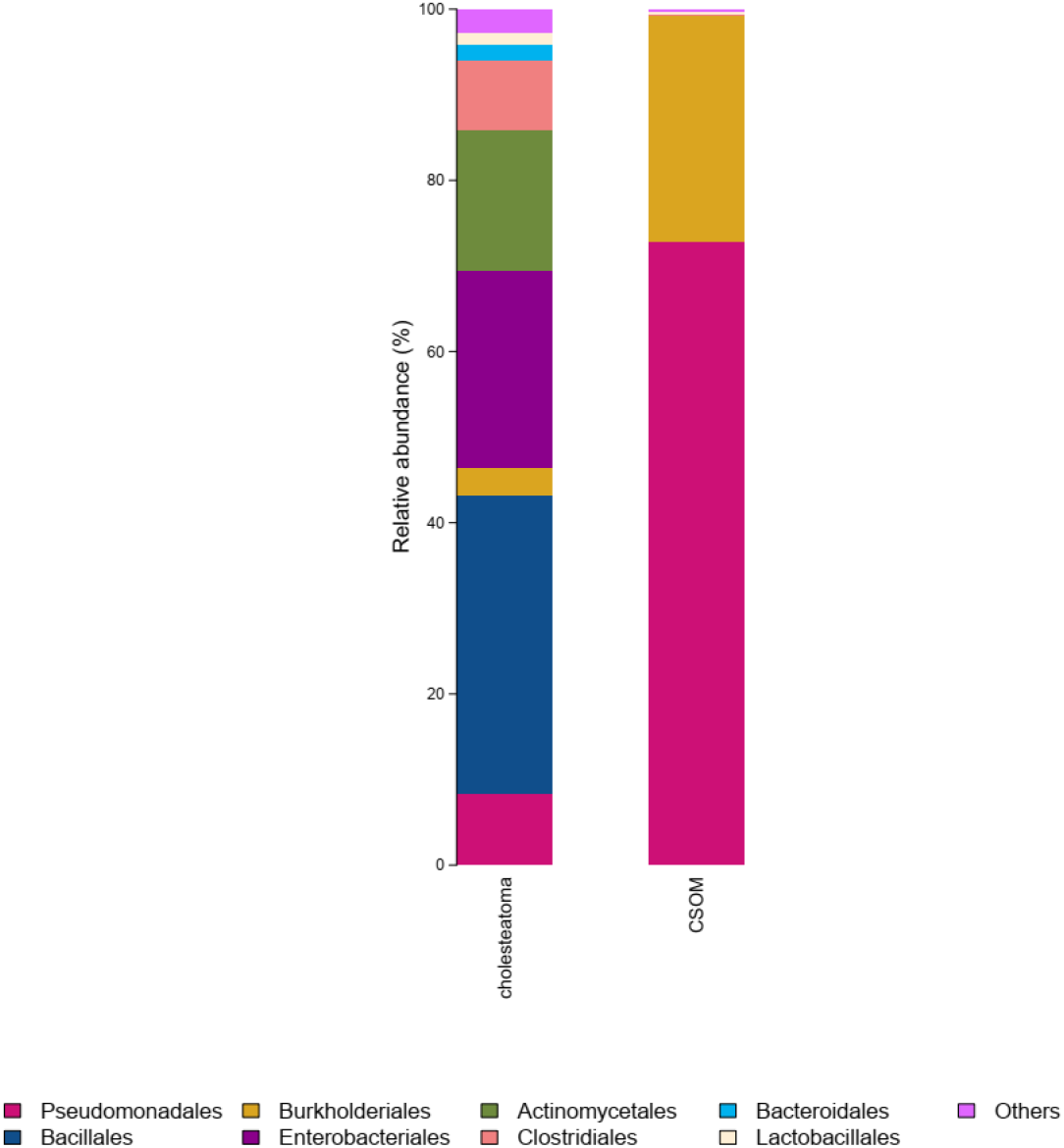
Bar plot diagram showing bacterial composition at the order level in CSOM and cholesteatomas pathological tissues

### FAMILY

Analysis of sequencing results at the family level indicated that *Pseudomonadaceae* (37.74%), *Moraxellaceae* (35.13%), and *Alcaligenaceae* (26.13%) were predominant in the CSOM samples. In particular, detection of *Pseudomonadaceae* (P=0.000976) was statistically significant. Bacterial diversity at the family level in the cholesteatoma group was more complex than that of the CSOM group. Families with broad coverage in the cholesteatoma group were *Staphylococcaceae* (33.53%), *Enterobacteriaceae* (22.93%), *Brevibacteriaceae* (8.49%), *Pseudomonadaceae* (8.39%), and *Corynebacteriaceae* (6.31%). The statistically significant families *Porphyromonadaceae* and *Lachnospiraceae* are mostly conditional pathogenic bacteria (FIG 8, FIG 9).

**FIG 8.**
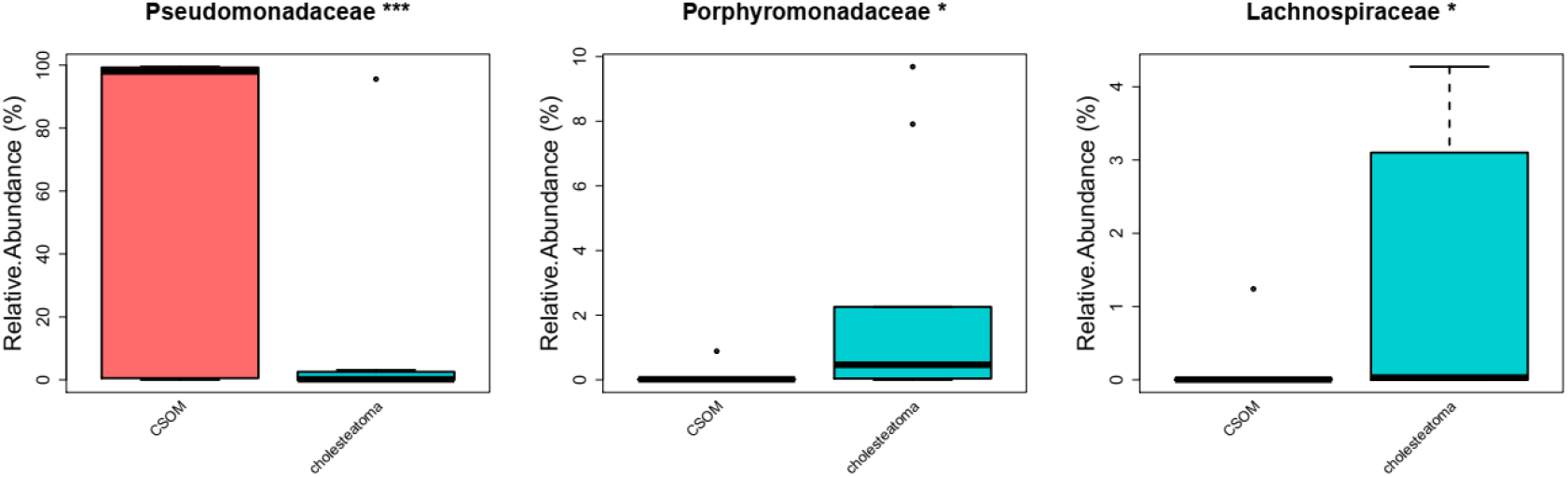
Family level differences in bacterial communities of CSOM and cholesteatoma lesions

**FIG 9.**
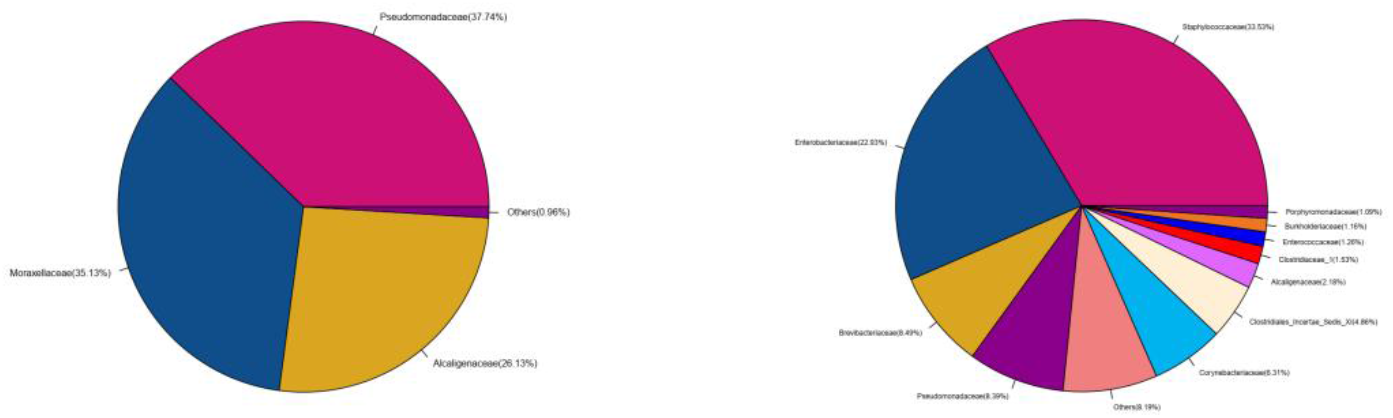
Pie charts showing composition of bacterial communities at the family level in CSOM and cholesteatoma lesions

### Genus

Sequencing analysis of bacterial communities of CSOM and cholesteatoma lesions at the genus level revealed that 16 genera, including ‘Others’ and ‘No_Rank’, were detected across the two groups. However, only *Pseudomonas* was detected in both groups and showed a statistically significance difference in relative abundance between the two groups (P=0.000601) (FIG 9). *Pseudomonas* in the CSOM group (37.74%) was significantly higher than that in the middle ear cholesteatoma group (8.39%). Other genera detected in the CSOM group were *Acinetobacter* (35.13%) and *Alkaligenes* (26.02%) (FIG 10). A total of the G-negative genera were dominant. These genera include *Pseudomonas aeruginosa*, *Acinetobacter baumannii*, and fecal alkaloid bacteria, which are the most common resistant strains of nosocomial infections. In contrast, the composition of bacterial genera of the cholesteatoma group was more elaborate, comprising S*taphylococcus* (33.53%), *Providencia* (12.34%), Others (9.56%), *Proteus* (9.05%), *Brevibacterium* (8.49%), *Pseudomonas* (8.39%), *Corynebacterium* (6.31%), *Finegoldia* (2.74%), No_Rank (2.15%), *Achromobacter* (2.09%), *Clostridium* (1.53%), *Tissierella* (1.32%), *Enterococcus* (1.26 %), and *Ralstonia* (1.08%) (FIG 10). These genera are diverse and include Gram-positive and Gram-negative bacteria, cocci and bacilli, aerobic and anaerobic bacteria, and so on.

**FIG 9.**
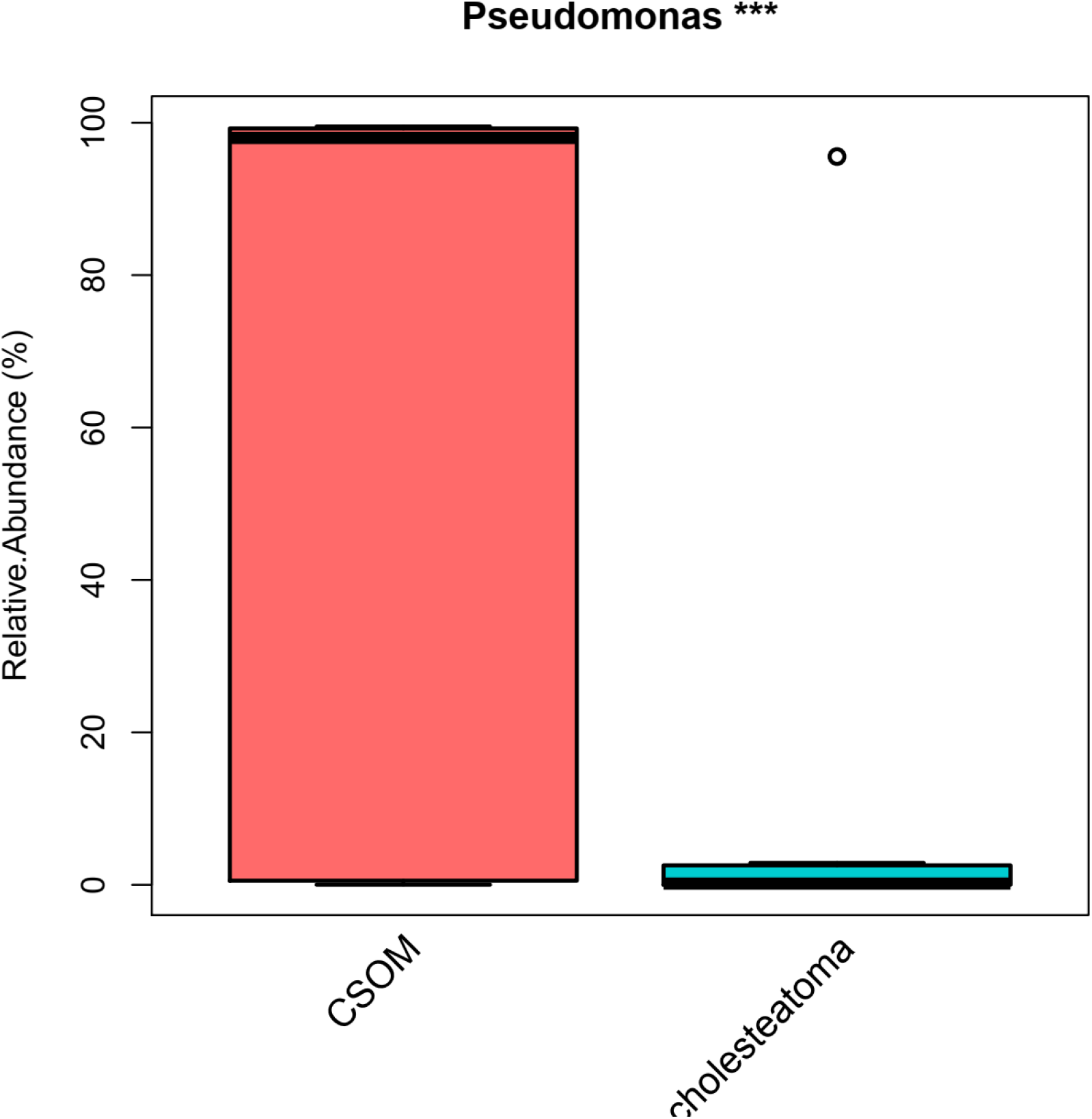
Genus level differences in bacterial communities of CSOM and cholesteatoma lesions

**FIG 10.**
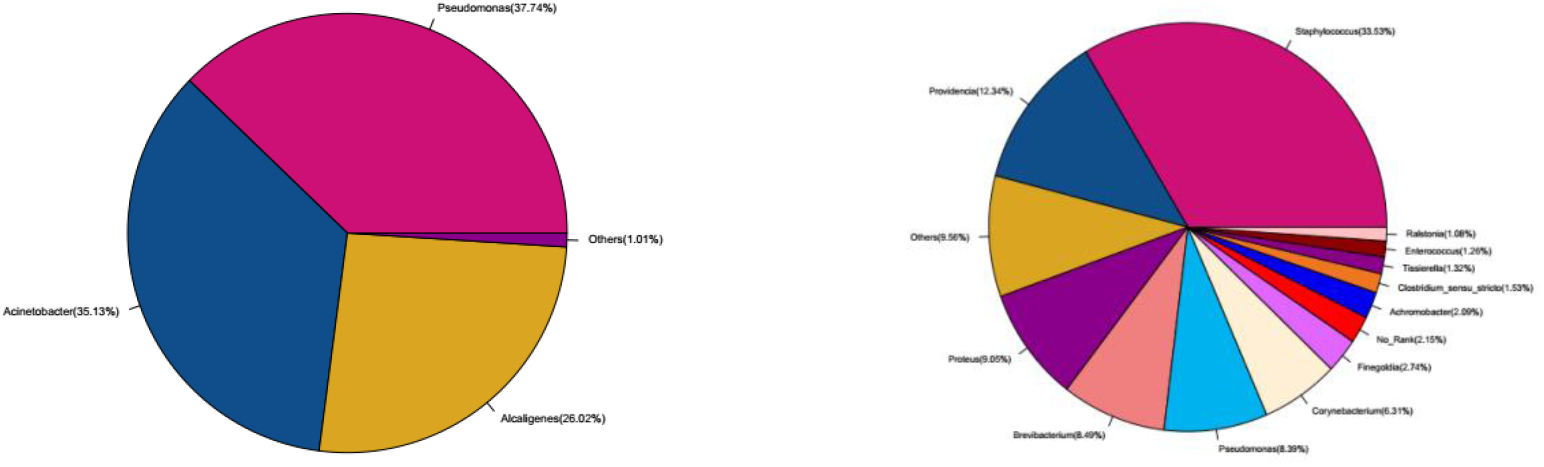
Pie charts showing genus level composition of bacterial communities in CSOM and cholesteatoma lesions

### Species

Sequencing analysis at the species level demonstrated that the composition of pathogenic bacterial species in cholesteatoma lesions was more complex than that of CSOM lesions. And the types of pathogenic bacteria were detected much more than CSOM. The statistical results is uncultured-organism (P=0. 001476), and uncultured-bacterium (P=0.0043290). The detection rate of *Pseudomonas aeruginosa* in the CSOM group was much higher than that in the middle ear cholesteatoma group (P=0. 000576) (FIG 11, FIG 12).

**FIG 11.**
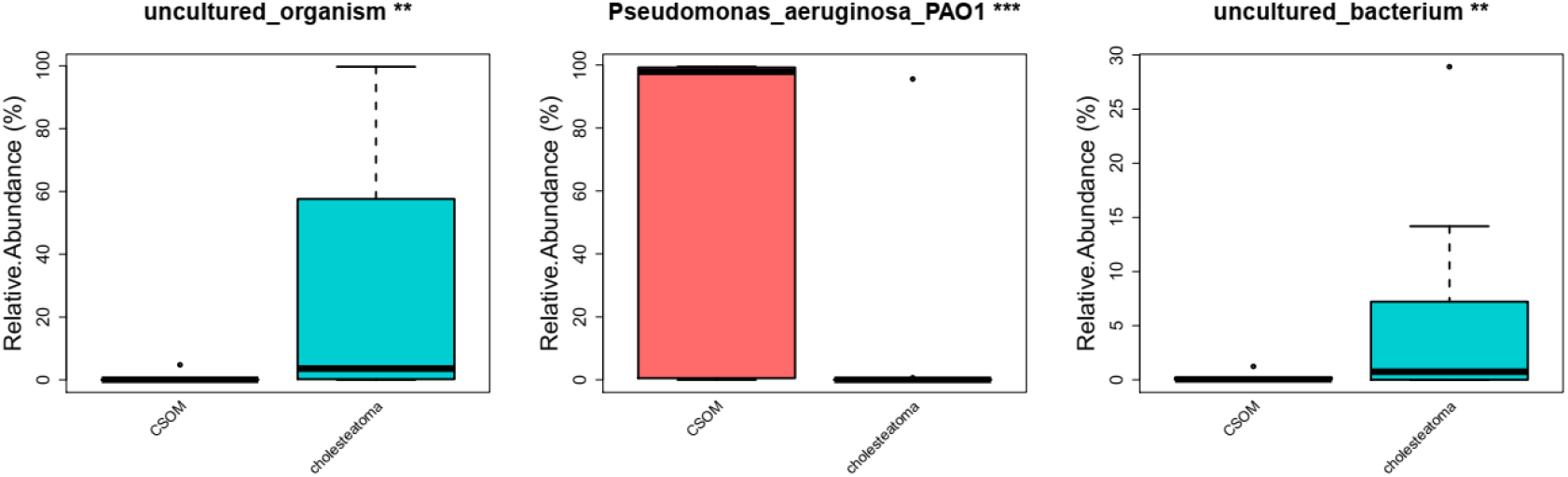
Species level differences between bacterial communities of CSOM and cholesteatoma lesions

**FIG 12.**
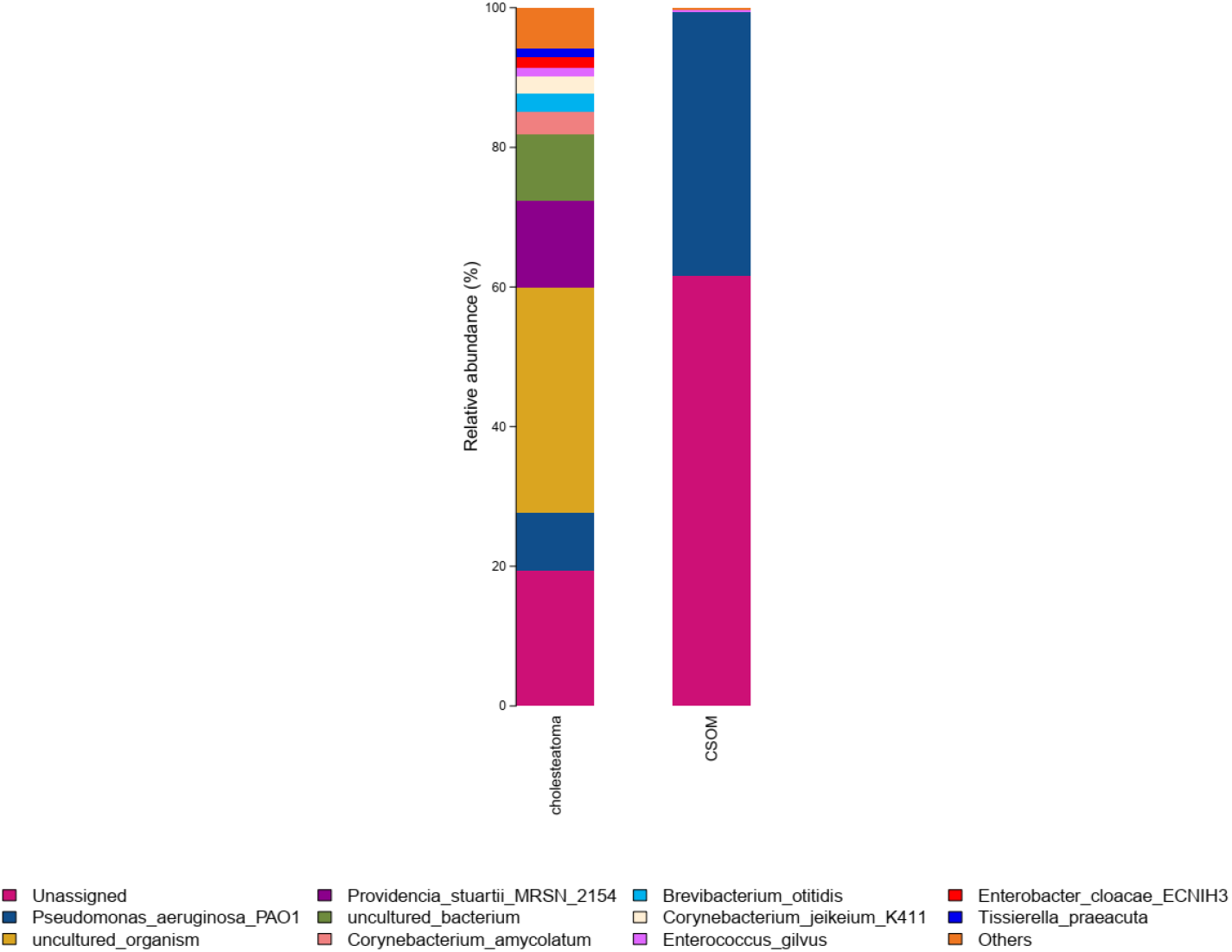
Bar plot showing species composition of bacterial communities of CSOM and cholesteatoma lesions

## DISCUSSION

Although the clinical manifestations of each type of otitis media are different, both CSOM and middle ear cholesteatoma require surgical treatment to cure the infection. There are limited differences in the clinical use of antibiotics for anti-infection treatment of two types of otitis media similar, and selection of an appropriate antimicrobial agent is solely based on ear secretion culture and drug sensitivity test. In practical clinical work, the results of bacterial culture of ear secretions often do not meet the purpose of curing otitis media (17). Clinicians seldom treat CSOM and middle ear cholesteatoma differently and do not select individualized treatments. Thus, there are limited studies on the detailed bacteriological differences between chronic otitis media and middle ear cholesteatoma.

In almost all molecular biological studies on the bacteriology of chronic otitis media, secretion swabs are collected from the middle ear cavity or the outer ear canal for gene sequencing. However, the collected secretions can be contaminated by bacteria in the external auditory canal or can be affected by the acquisition process. This may lead to false positive results and thus these studies do not accurately reflect the true bacteriology of chronic otitis media (12,18,19). Simultaneously, the formation of bacterial biofilm, like in many chronic inflammatory diseases, severely limits the ability to obtain positive results by conventional bacterial culture. Consequently, these analyses on chronic otitis media do not reflect the true clinical situation (20,21).

An alternative approach, utilizing gene sequencing to analyze unusual or dominant flora in the diseased tissue, may be beneficial for the study of the pathogenesis of otitis media. In this prospective study, high-throughput gene sequencing of diseased tissues of CSOM and cholesteatoma was performed). Bacterial species abundance and species diversity both differed in the two types of otitis media. Bacterial species abundance and diversity of the cholesteatoma group was greater than that of the CSOM group. This finding is congruent with previous studies (22). Collectively data from these studies suggest that the bacterial infection situation in middle ear cholesteatoma may be different and more complex than that of CSOM.

The current study revealed that the bacterial community composition in the middle ear lesion tissue of cholesteatoma was significantly more complex than that of CSOM. Lesions of the cholesteatoma group contained numerous Gram-positive and Gram-negative bacteria, which may be related to the patient’s long-term illness, with the bacteria forming a unique microenvironment, a coexistence relationship, and biofilm. In contrast, the bacteriology of CSOM lesions is relatively simple, with bacterial communities at all taxonomic levels (phyla, classes, orders, families, genera, and species) dominated by Gram-negative bacteria. These results indirectly explain the clinical phenomenon that broad-spectrum antibiotics are sometimes effective in patients with CSOM but have no effect on patients with middle ear cholesteatoma.

The findings of the current study differ from previous bacteriological studies on chronic otitis media which usually only produce one or two kinds of bacteria. This is due to difficulties in cultivating a variety of bacteria using conventional bacteriological techniques, meaning such studies cannot reflect the full picture of bacterial infection in patients. However, using high-throughput gene sequencing, the detection rate of pathogenic bacteria in middle ear cholesteatoma lesions was significantly higher than that in CSOM lesions. This indicates that cholesteatoma has a much higher complexity of bacterial species and it is not an infectious disease caused by a single pathogenic bacterial species. Cholesteatoma may therefore constitute a bacterial microenvironment, whereby the combination of multiple conditional pathogens leads to the continuous onset of clinical symptoms of middle ear cholesteatoma, and a series of complications eventually emerges. This finding warrants further attention in future research.

Based on results of the current study, it is suggested that in future clinical work, antimicrobial agents are rationally selected according to the disease types of patients in addition to the clinical microbial culture and drug sensitivity test. Patients with CSOM could use broad-spectrum antibiotics against Gram-negative bacteria, but relying solely on antibiotics would most likely be ineffective for middle ear cholesteatoma. After surgical removal of diseased tissue, appropriate antimicrobial agents against infection could be selected in combination with actual drug sensitivity tests. Furthermore, considering the complexity of the bacterial infection would avoid drug resistance and flora disorders caused by the excessive use of broad-spectrum antimicrobial therapy.

## ACKNOWLEDGMENTS

We thank all participants of this study. Colleagues of the otolaryngology diagnosis and treatment center of Xinjiang Uygur Autonomous Region People’s Hospital are acknowledged for their hard work in collecting cases and specimens.

## FUNDING INFORMATION

The author(s) disclosed receipt of the following financial support for the research, authorship, and/or publication of this article: This study was financially supported by the National Science Foundation of China, Project No. 81560173.

